# The genetic architecture of resistance to flubendiamide insecticides in *Helicoverpa armigera* (Hübner) (Lepidoptera: Noctuidae)

**DOI:** 10.1101/2024.07.28.605483

**Authors:** Douglas Amado, Eva L. Koch, Erick M. G. Cordeiro, Wellingson A. Araújo, Antonio A. F. Garcia, David G. Heckel, Gabriela Montejo-Kovacevich, Henry L. North, Alberto S. Corrêa, Chris D. Jiggins, Celso Omoto

## Abstract

Insecticide resistance is a major problem in food production, environmental sustainability, and human health. The cotton bollworm *Helicoverpa armigera* is a globally distributed crop pest affecting over 300 crop species. *H. armigera* has rapidly evolved insecticide resistance, making it one of the most damaging pests worldwide. Understanding the genetic basis of insecticide resistance provides insights to develop tools, such as molecular markers, that can be used to slow or prevent the evolution of resistance. We explore the genetic architecture of *H. armigera* resistance to a widely used insecticide, flubendiamide, using two complementary approaches: genome-wide association studies (GWAS) in wild-caught samples and quantitative trait locus (QTL) mapping in a controlled cross of susceptible and resistant laboratory strains. Both approaches identified one locus on chromosome 2, revealing two SNPs within 976 bp that can be used to monitor field resistance to flubendiamide. This was the only region identified using linkage mapping, though GWAS revealed additional sites associated with resistance. Other loci identified by GWAS in field populations contained known insecticide detoxification genes from the *ATP-binding cassette* family, ABCA1, ABCA3, ABCF2 and MDR1. Our findings revealed an oligogenic genetic architecture, in contrast to previous reports of monogenic resistance associated with the *ryanodine receptor*. This work elucidates the genetic basis of rapidly evolving insecticide resistance and will contribute to the development of effective insecticide resistance management strategies.

**Author summary:** Insecticide resistance in agricultural pests challenges food security, environmental sustainability, and human health. The cotton bollworm *Helicoverpa armigera* is resistant to various insecticide classes as well as *Bacillus thuringiensis* toxins. Understanding the genetic basis of this resistance is crucial for developing effective insecticide resistance management (IRM) strategies. Our study investigated the genetic architecture of resistance of *H. armigera* to flubendiamide and identified SNPs associated with resistance. We used two approaches: association mapping in wild-derived samples and QTL mapping in a controlled backcross of susceptible and resistant laboratory strains. One specific region on chromosome 2 was found in both GWAS and QTL mapping analysis and showed significant potential as a PCR-based marker for monitoring resistance to flubendiamide in *H. armigera*. Additionally, the GWAS identified five more SNPs in the field populations. The candidate genes identified are primarily associated with insecticide detoxification mechanisms and calcium homeostasis. Our study advances the understanding of the complex genetic architecture of flubendiamide resistance in *H. armigera,* providing valuable tools for IRM programs and establishing a basis for future studies.

## Introduction

Insecticides play an important role in agricultural pest management and controlling vectors of human diseases. However, their frequent application has led to the rapid evolution of resistant populations, representing challenges to pest management efforts. The development of insecticide resistance threatens sustainability and human health due to the increased frequency and rate of insecticide applications [1,2]. Insecticide Resistance Management (IRM) programs are necessary for creating sustainable and long-term techniques to mitigate the evolution of resistance in pest species. A fundamental component of these programs is understanding the genetic basis of insecticide resistance and monitoring allele frequencies within field populations before control failures occur [3–5].

Insecticide resistance in insects is affected by genetic variation in the population, mutational constraints on genes, and the selection pressure imposed by insecticide use [6]. These factors can lead to monogenic, polygenic, and oligogenic insecticide resistance. Monogenic resistance typically results from a single, rare mutation with a major effect, often identified within target-site genes. When the individuals of a field population are exposed to a high dose of insecticide, the mutations with a major effect may be selected, leading to a rapid increase in the resistance allele frequency [7]. Conversely, polygenic resistance is caused by many mutations with minor effects. In the absence of alleles with major effects, the individuals of a field population systematically exposed to a low dose of insecticide over several generations can accumulate mutations with minor or intermediate effects, leading to a slow increase in resistance allele frequency [3,8]. Between these two extremes lies oligogenic resistance, in which a few mutations with major effects account for most of the trait variation, along with minor/intermediate-effect mutations that modify the resistance phenotype [9,10]. In the field, populations can be exposed to different insecticides in multiple applications across time, and the selection of mutations with major and minor effects makes it difficult to predict insecticide resistance evolution [10].

In this context, traditional laboratory mortality assays used to screen for resistance often lack the sensitivity to monitor and discriminate the major and minor alleles related to insecticide resistance in the field, especially when these alleles are at low allele frequencies [4]. Furthermore, laboratory-selected populations often used to study resistance can exhibit polygenic resistance mechanisms that do not represent those seen in the field [11].

An alternative approach is to use multiple molecular markers to monitor allele frequencies in pest populations [5,12,13]. Both Genome-Wide Association Studies (GWAS) and Quantitative Trait Loci (QTL) mapping have been used to identify markers associated with resistance, each with distinct advantages and limitations [14–18]. Combining GWAS and QTL mapping together enhances the ability to study the genetic architecture and identify and refine alleles linked to interest traits. While QTL mapping can identify loci that control a trait and estimate their effect size and genetic × environmental interactions, GWAS can narrow down candidate regions, detecting minor or rare alleles [4,19,20]. Thus, GWAS results can be validated through QTL mapping in crossed populations, and conversely, the QTLs identified can be examined in natural populations by GWAS [4,20,21]. Few studies have applied this combined approach to the complex genetic architecture of insecticide resistance in agriculture pests.

The insecticide flubendiamide comprises the group of diamides, more specifically, the phytalic acids sub-group. This insecticide was commercially released in 2007, with a different mechanism of action compared to existing products, effectiveness against lepidopteran and coleopteran pests, and low toxicity to mammals [22]. Flubendiamide acts by irreversibly binding to *ryanodine receptors* (RyRs) in the sarco/endoplasmic reticulum, causing uncontrolled Ca^2+^ efflux. This calcium acts on muscle contraction, but the excess of calcium promotes muscle paralysis, leading to the insect’s death [23–26]. However, the extensive application of this insecticide has led to control failures worldwide due to an increase in resistance evolution [27]. Resistance to flubendiamide has been reported in some species, including *Plutella xylostella* (L.) (Lepidoptera: Plutellidae), *Spodoptera exigua* (Hübner) (Lepidoptera: Noctuidae), *Tuta absoluta* (Meyrick) (Lepidoptera: Gelenchiidae), and *Spodoptera frugiperda* (Smith) (Lepidoptera: Noctuidae) [18,28–32]. By monitoring the susceptibility of field populations of *Helicoverpa armigera* (Hübner) (Lepidoptera: Noctuidae) during the 2014 to 2018 crop seasons, it was possible to verify a reduction of flubendiamide susceptibility in different regions of Brazil [33]. In this way, an *H. armigera* laboratory strain was established in the 2016 crop season, which reached resistance ratios exceeding 50,000-fold to flubendiamide after being selected in the lab for a couple of generations [34].

Since the release of flubendiamide, many studies have focused on understanding the genetic mechanisms of flubendiamide resistance. These resistance mechanisms are linked to specific mutations in the T2 to T6 transmembrane domains of the C-terminal region of RyR and with differential expression of detoxication enzymes such as *cytochromes P450* (P450), *carboxylesterases* (CE), and *glutathione S-transferases* (GST) [18,35–42]. In contrast, the *H. armigera* laboratory-selected strain (> 50,000-fold) exhibits only synonymous mutations in the RyR and no alterations in mortality associated with insecticide synergists such as P450, CE, and GST [34]. Thus, there is a gap in knowledge of the genetic architecture of flubendiamide resistance in *H. armigera* in Brazil. Unlike previous works, we employed a more robust and comprehensive approach by combining GWAS and QTL mapping studies to investigate the genetic architecture of the flubendiamide resistance in *H. armigera*. This information is crucial for understanding the evolutionary process associated with insecticide resistance and its management in the field, especially given the rapid evolution of flubendiamide resistance in *H. armigera* in Brazil.

## Results

### Genome-Wide Association Studies (GWAS)

We used the GBS from 110 individuals from a field population of *H. armigera* to conduct a GWAS for flubendiamide resistance. After data filtering, 103 individuals and 9,259 SNPs were retained for analysis. Of these individuals, 47 (46%) were observed to be sensitive to flubendiamide, while 56 (54%) survived. Structure analysis sorted these 103 individuals into three clusters (K = 3), as shown in (S1 Fig). The number of clusters was incorporated into the association model to minimize the potential for false positives. Linkage disequilibrium (LD) decay analysis suggested a window size of approximately 10 Kb. However, we also employed a 300 Kb window, as suggested by Anderson et al. (2018) [43]. Employing the Blink model for the genotype-phenotype association, we found a good fit for the data (S2 Fig). Four markers showed significant associations, surpassing the Bonferroni-corrected threshold (*p*-value = 5.4×10^-6^) (Fig 1 and Table 1).

**Fig 1.**
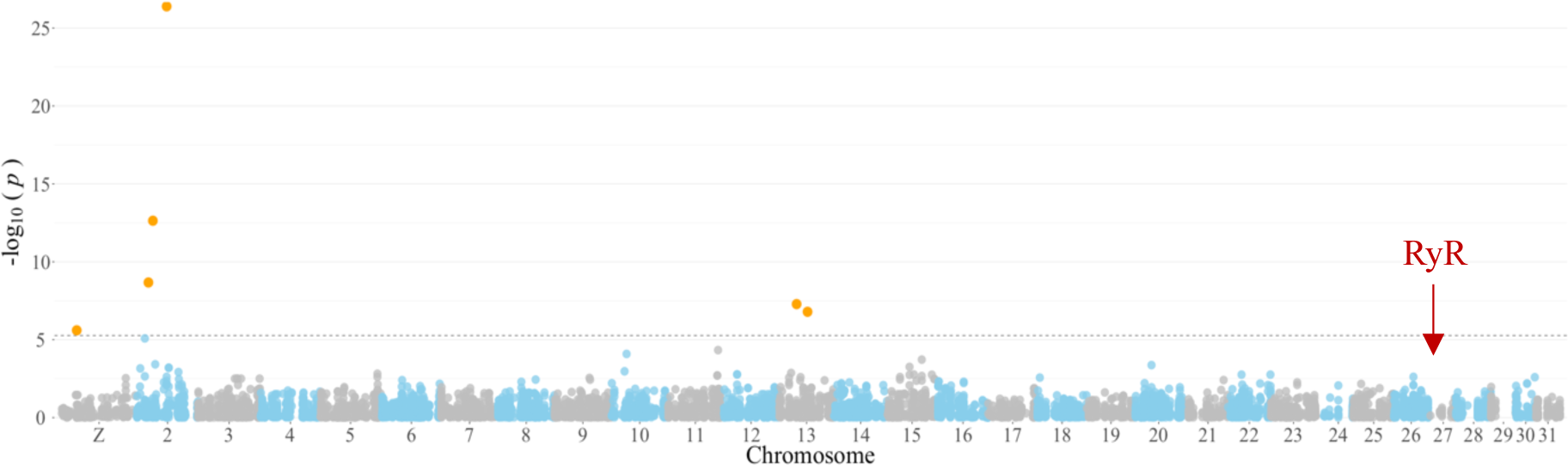
Genome-wide association plot of single nucleotide polymorphisms associated with flubendiamide survival traits in field-derived *Helicoverpa armigera*. The x-axis denotes the number and position of markers across chromosomes. The y-axis illustrates the −log10(*P-values*) to depict the significance of each SNP. A horizontal dashed line indicates the Bonferroni-adjusted significance threshold of 5.4×10^-6^. A red arrow identifies the location of the ryanodine receptor gene on the *H. armigera* reference genome.

**Table 1.** Markers identified in the genome-wide association of field-collected *H. armigera* individuals associated with flubendiamide survival trait.

| Marker | RefSeq<sup>a</sup> | Chr | Position | *p*-value | Maf<sup>b</sup> | Effect | PVE<sup>c</sup> |
| --- | --- | --- | --- | --- | --- | --- | --- |
| rs1P3460302 | NC_087120.1 | Z | 3460302 | $2.494 \times 10^{-06}$ | 0.131 | 0.150 | 2.352 |
| rs2P2759433 | NC_087121.1 | 2 | 2759433 | $2.133 \times 10^{-09}$ | 0.116 | -0.285 | 8.388 |
| rs2P3779183 | NC_087121.1 | 2 | 3779183 | $2.318 \times 10^{-13}$ | 0.155 | 0.392 | 10.382 |
| rs2P6931787 | NC_087121.1 | 2 | 6931787 | $4.034 \times 10^{-27}$ | 0.106 | 0.904 | 27.774 |
| rs13P4171996 | NC_087132.1 | 13 | 4171996 | $5.197 \times 10^{-08}$ | 0.024 | 0.424 | 20.865 |
| rs13P6678940 | NC_087132.1 | 13 | 6678940 | $1.622 \times 10^{-07}$ | 0.043 | 0.429 | 6.926 |

<sup>a</sup>NCBI genome reference sequence assembly (GCF\_030705265.1).
<sup>b</sup>Minor allele frequency.
<sup>c</sup>Percentage of phenotypic variation explained by the marker.

Among the six significant markers, the markers rs2P2759433, rs2P3779183, and rs2P6931787, located on chromosome 2, accounted for 8%, 10%, and 27% of the phenotypic variation, respectively (Table 1). One additional marker on chromosome Z and two markers on chromosome 13 were collectively responsible for 30% of the variation (Table 1). All six markers explained approximately 77% of the observed survival phenotype.

### QTL mapping

Next, we performed phenotype and genotype analyses on a sample of 109 individuals from a backcross population derived from the cross between susceptible and resistant strains of *H. armigera* to flubendiamide. Among these, 49 individuals (45%) were susceptible to the diagnostic dose of flubendiamide, while 60 (55%) exhibited resistance. We developed a linkage map for *H. armigera* for the 31 linkage groups using 1,118 markers that met our stringent criteria (S3 Fig). These groups varied in size from 91.56 to 153.25 cM, with a total length of 3,722.75 cM (S1 Table).

The genotype proportions across the linkage groups were 50.4% for AA and 49.6% for AB. Our QTL mapping identified a single significant QTL associated with flubendiamide resistance on chromosome 2, which had an LOD score of 6.92, surpassing the threshold of 3.30 established by permutation tests (Fig 2). This QTL accounts for 25.5% of the phenotypic variation. At the peak of this QTL, the SNP, rs2P2760409 was located, and the upper confidence interval was identified the SNP rs2P14104337 in the position 8.31 cM (physical position 14,104,337 bp) (Table 2). Given the relatively small proportion of variance explained, the QTL result is also consistent with resistance to flubendiamide being an oligogenic trait, as the identified QTL does not fully account for the resistance phenotype.

**Fig 2.**
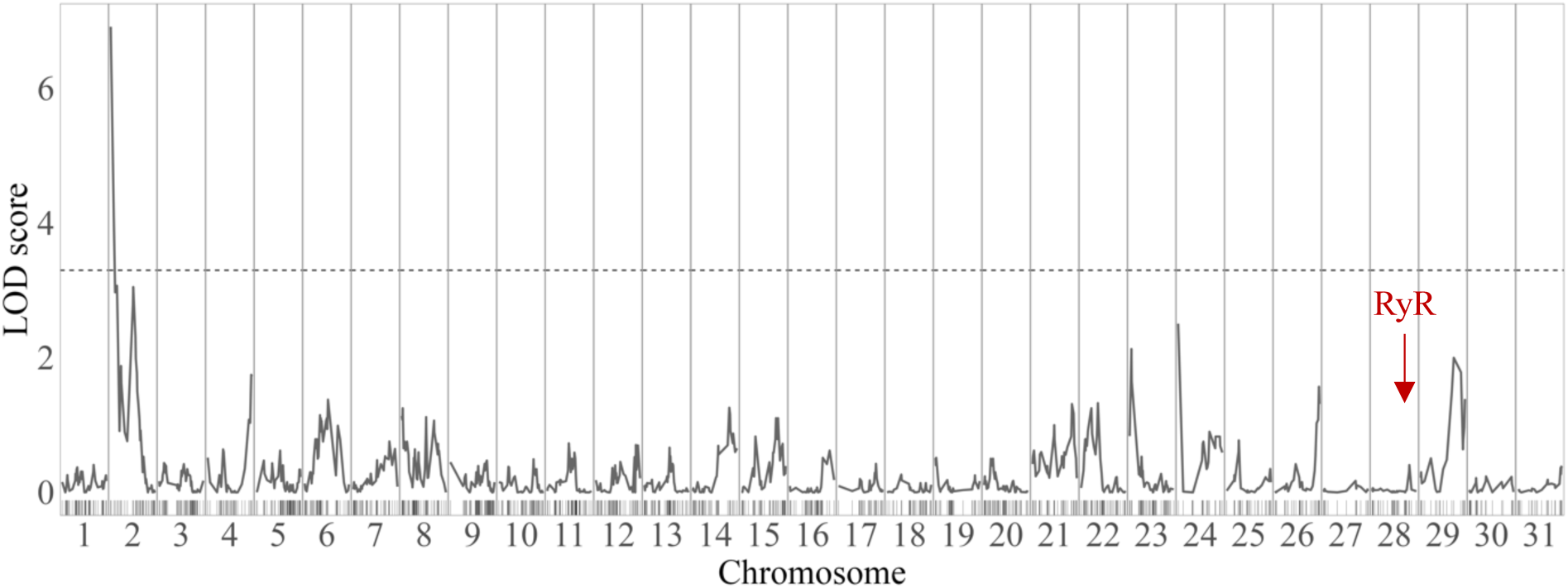
QTL mapping for flubendiamide resistance in the *Helicoverpa armigera* Flub-R laboratory strain. The x-axis details the chromosomes and marker positions, while the y-axis displays the LOD scores. A horizontal dashed line marks the significance threshold at an LOD of 3.30, as determined by permutation tests. A red arrow highlights the location of the ryanodine receptor gene on the *H. armigera* reference genome.

**Table 2.** Localisation and LOD scores of markers within QTL peak intervals for flubendiamide resistance on linkage group 2 in the *Helicoverpa armigera* Flub-R laboratory strain.

| SNP ID | Linkage Group | RefSeq<sup>a</sup> | Position (cM)<sup>b</sup> | LOD |
| --- | --- | --- | --- | --- |
| rs2P2760409 | 2 | NC_087121.1 | 6×10<sup>-6</sup> | 6.917 |
| rs2P1937820 | 2 | NC_087121.1 | 0.918 | 6.550 |
| rs2P14104337 | 2 | NC_087121.1 | 8.310 | 3.863 |

<sup>a</sup>NCBI genome reference sequence assembly (GCF\_030705265.1).
<sup>b</sup>Genetic position of the SNP on linkage group (in centiMorgans).

Therefore, the QTL mapping and GWAS results are highly consistent. In terms of physical position, the SNP rs2P2759433 identified by GWAS is approximately 976 bp from the SNP rs2P2760409 in the QTL peak (Fig 3). This region, therefore, plays a significant role in the phenotype of *H. armigera* resistance to the insecticide flubendiamide.

**Fig 3.**
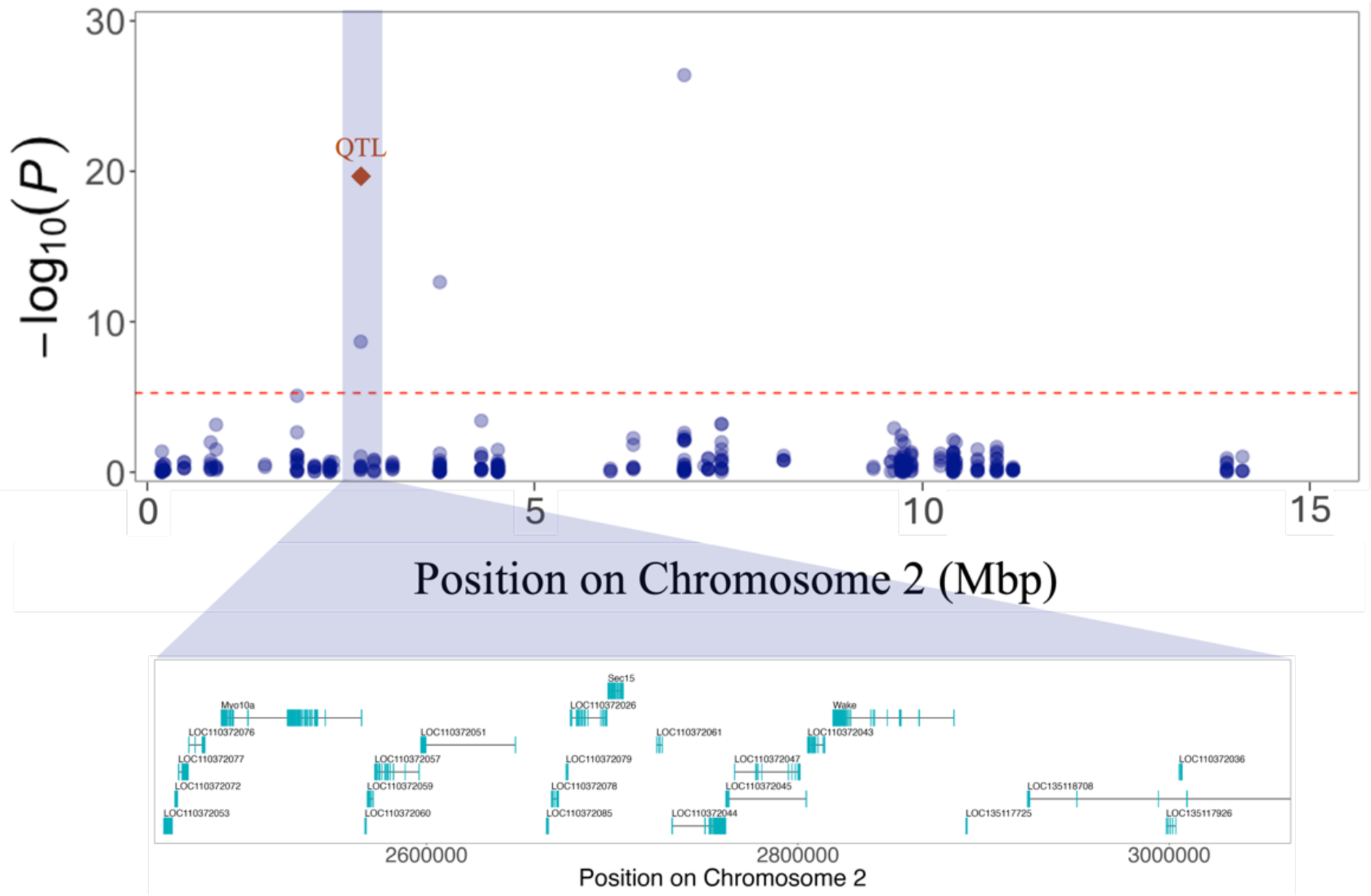
Overlap of GWAS and QTL mapping results. The x-axis represents the physical position of the SNPs, while the y-axis shows the *p-value* of the GWAS result transformed by −log10(*p*-values). The red dashed line represents the Bonferroni-corrected GWAS threshold (*p-value* = 5.4×10^-6^). The blue dots represent the SNPs from the GWAS, while the orange diamond indicates the physical position of the SNP identified at the QTL peak from the QTL mapping analysis. The lower panel displays the genes identified within a 300 kb window upstream and downstream of the SNP rs2P2759433 identified by GWAS and the SNP rs2P2760409 identified by QTL Mapping.

### Candidate genes

Identification of specific candidate genes is necessarily speculative at this stage due to broad confidence intervals around QTL and GWAS SNPs. However, in order to investigate possible candidates, regions spanning 300 kb upstream and downstream of SNPs identified by GWAS and QTL analyses were scanned for possible candidate genes. In the window surrounding the SNP rs1P3460302 on chromosome 1, 16 characterised genes were found. Among these are the *Calcium-binding Mitochondrial Carrier Protein SCaMC-2* and *ATP-binding Cassette Subfamily F Member 2* genes (S2 Table). Furthermore, the *Developmentally-regulated GTP-binding Protein 2* gene was identified adjacent to SNP rs2P2759433 and rs2P2760409. These SNPs were co-located by GWAS and QTL Mapping analyses (Fig 3). In the region of SNP rs2P3779183, 30 characterised genes were identified, including the *Voltage-dependent T-type Calcium Channel Subunit Alpha-1G* gene. In the same way, 38 characterised genes were found in the window of SNP rs2P6931787, of which the *Phospholipid-transporting ATPase ABCA1* and *Phospholipid-transporting ATPase ABCA3* are strong candidates for the high survival of field population (S2 Table). Of the 36 characterised genes identified within the SNP window of rs13P4171996, the *Multidrug Resistance-associated Protein 1* gene is the most likely to be associated with the high survival rate of the field population. Similarly, the *Cytochrome C Oxidase Assembly Protein COX20* gene, identified among the 18 characterised genes within the SNP region of rs13P6678940, is also a strong candidate. Both markers are located on chromosome 13 (S2 Table).

## Discussion

We have identified a single major-effect locus influencing resistance to flubendiamide in Brazilian *H. armigera*, supported by independent QTL and GWAS analyses. In addition, there is evidence for five additional loci with more minor phenotypic effects. Previous studies have identified monogenic inheritance of diamide resistance in agricultural pests, mainly associated with mutations in the T2 to T6 transmembrane domains of the C-terminal region of RyR [18,28,36,37,39,40]. Our previous work characterising resistance of the Flub-R strain to the insecticide flubendiamide did not reveal any non-synonymous mutations in the *RyR*, nor did it show any changes in the mortality of individuals exposed to flubendiamide combined with the synergists PBO, DEM, and DEF [34]. This absence suggests the involvement of an alternative resistance mechanism in *H*. *armigera*.

We confirm this result here, as our GWAS and QTL Mapping did not identify any associations on chromosome 28 associated with *RyR*. We have identified a new resistance locus and highlighted the contribution of additional minor effect genes influencing the survival of *H. armigera* to flubendiamide. [44]

The GWAS analysis identified genetic loci with a major-effect on chromosome 2 and loci with a minor effect on chromosomes 1 and 13. Together, these loci are responsible for much, but not all, phenotypic variation. These loci had high statistical support and somewhat overcame the limitations of traditional laboratory assays with field populations, which may not effectively estimate the minor-effect alleles found in field populations [4,13,45]. Using a GWAS approach with a field population increased the probability of the allele(s), with the major-effect being sampled and identified [46].

In parallel, we identified one major-effect QTL associated with *H. armigera* resistance to flubendiamide from a laboratory-selected strain. This QTL on chromosome 2 explained part of the phenotypic variance, showing that the flubendiamide-resistant strain (Flub-R) has an oligogenic genetic architecture, underscoring the complexity of flubendiamide resistance. This contrasts with previous studies, which have demonstrated monogenic resistance, often linked to mutations in *RyR* or differential expression of detoxification enzymes [36]. The statistical significance of the QTL and narrow confidence intervals supports the robustness of our findings.

Furthermore, the concordance between GWAS and QTL analysis in the identification of the same locus on chromosome 2 provides strong support for the importance of this genomic region for *H. armigera* resistance to flubendiamide. The SNPs rs2P2759433 and rs2P2760409 are promising for developing PCR-based markers to complement traditional phenotypic laboratory assays in resistance monitoring. This should significantly enhance the sensitivity and accuracy of these assessments [12,13], offering a more refined strategy for resistance management. The combined use of GWAS and QTL mapping has been extensively employed in studies of complex phenotypic traits in plant breeding [20,47–49]. Still, it has been less commonly used in insect resistance research. Our work demonstrated that this is a powerful approach to studying complex traits in insect populations, such as insecticide resistance in crop pests.

The genes near the SNPs co-located in the GWAS and QTL mapping imply a different resistance mechanism to flubendiamide. The *developmentally-regulated GTP-binding protein 2* genes identified in that region may be associated with insect resistance by insecticide detoxification. This association was observed in a differential expression study involving *Mythimna separata* (Walker) (Lepidoptera: Noctuidae) treated with chlorantraniliprole [50]. On the other hand, the *Voltage-dependent T-type Calcium Channel Subunit Alpha-1G* close to the SNPs rs2P3779183 on chromosome 2, identified only by GWAS analysis, is associated with the calcium homeostasis pathways [51,52].

Genes related to metabolic detoxification were observed in the region of the higher peak SNP (rs2P6931787) on chromosome 2, identified only by GWAS analysis. In this region, we found four possible *ABC transporter* genes that act in phase III of metabolic detoxification. Many studies have reported the involvement of *ABC transporters* in chlorantraniliprole and flubendiamide detoxification in other species such as *Chilo suppressalis* (Walker) (Lepidoptera: Crambidae) [53], *P. xylostella* [54], *S. frugiperda* [55] and *Leptinotarsa decemlineata* (Say) (Coleoptera: Chrysomelidae) [44]. The *Multidrug resistance-associated protein 1* (MDR1), also known as *P-glycoprotein* (P-gp), is encoded by *ATP-binding cassette subfamily B member 1* (ABCB1) [56,57]. P-gp, a component of the *ABC superfamily*, is a membrane-spanning protein that pumps molecules out of the cell through an ATP-dependent mechanism [58,59].

Studies have shown that some insecticides act as substrates, increasing the P-gp expression and enhancing *ABC transporter* activity. This enables the insecticide to be transported out of the cell, contributing to resistance [56,60–63]. The study by Zuo et al. (2017) involving P-gp knockout in *S. exigua* showed no increases in chlorantraniliprole susceptibility [61]. Thus, the expression of P-gp might be more closely associated with phthalic insecticides than with anthranilic insecticides. This could explain the difference observed between the insecticides chlorantraniliprole and flubendiamide susceptibility in field populations from the state of Bahia in our previous study [33].

In summary, our results indicate that Flub-R strain resistance to flubendiamide is oligogenic and that a major effect locus on chromosome 2 contributes to resistance and could be the target for genetic monitoring of resistance in field populations. Furthermore, genes from the *ATP-binding cassette* family may also play significant roles in *H. armigera* resistance to flubendiamide insecticide. However, it is important to note that the study was limited to a single field population and that functional validation of the identified genes will be necessary. Future research should focus on validating these genes and assessing their frequency in different field populations. These findings contribute to the development of more effective IRM strategies, helping mitigate the devastating ecological and economic impacts of insecticide resistance.

## Materials and methods

### Field population for GWAS analysis

One thousand individuals of *H. armigera* were collected in the field from a soybean crop in the municipality of Luiz Eduardo Magalhães, Bahia (12°05’58”S and 45°47’54’’W) during the 2019 crop season. This population was subsequently maintained in the laboratory up to the adult stage on a modified artificial diet [64]. On average, 300 adults were reared in each PVC cage covered at the top with fabric as an oviposition substrate. The fabric with eggs was replaced every two days. The newly hatched larvae (F1 generation) were then transferred into 100 mL plastic cups containing an artificial diet. These larvae were maintained under controlled environmental conditions at 25±1°C, with relative humidity (RH) of 70±10%, and a 14:10 h light/dark photoperiod. The beginning of F1 third instar larvae were used for phenotypic bioassay. The F1 larvae were used for the phenotypic bioassay due to the difficulty of standardizing the size of individuals collected in the field.

### Segregating backcross population for QTL mapping

The *H. armigera* flubendiamide-resistant strain (Flub-R) originated from individuals collected on soybean in Luiz Eduardo Magalhães, Bahia, Brazil (12°05’58’’S and 45°47’54’’W) in the 2016 crop season. These individuals were initially selected in the susceptibility monitoring using a diagnostic dose (2.64 μg a.i. cm^-2^) of flubendiamide (Belt^®^, Bayer S.A.; 480 g a.i. L^-1^) and maintained under insecticide selection for several generations. The Flub-R strain exhibited a resistance ratio of more than 50,000-fold, as reported by Abbade-Neto et al. [34]. The susceptible *H. armigera* strain (TWBS) is from Australia and has been maintained without insecticide selection in the laboratory for more than 40 generations.

For QTL mapping, reciprocal couples were established with the Flub-R and TWBS strains (♀ TWBS × ♂ Flub-R and ♀ Flub-R × ♂ TWBS). The heterozygote individuals (HET) generated by the couple ♀ TWBS x ♂ Flub-R were used for backcrossing with individuals of the susceptible strain (TWBS). In the same way, from the HET and TWBS strains were established reciprocal couples (♀ TWBS × ♂ HET and ♀ HET × ♂ TWBS) to originate the backcross population (S4 Fig). The resulting offspring, segregating backcross 1 (BC1) from the couple ♀ TWBS × ♂ HET, was submitted to phenotypic bioassays.

### Phenotypic bioassays

Phenotyping of *H. armigera* larvae for QTL mapping and GWAS analysis was conducted using a dose-response bioassay. This involved the superficial application of the artificial diet with 30 µL of a diagnostic dose of 2.64 μg, a.i. cm^-2^ of flubendiamide (LD_99_) per cell [33]. The dose was prepared in distilled water with 0.1% Triton^®^ X-100 surfactant. For the bioassay, 24-cell acrylic plates, each cell containing 1.25 mL of diet, were used. After drying the insecticide solution in a laminar flow chamber, early third-instar larvae were individually placed in each cell. A total of 110 larvae from the field population were tested for GWAS, and 110 larvae from BC1 were tested for QTL, with each larva representing a single repetition. The plates were maintained at 25±2°C under a 14:10 h light/dark photoperiod. Mortality was assessed after 96 hours, with immobile individuals upon prodding considered dead. Phenotypic data were coded with *’1’* for survival and *’0’* for dead.

### DNA extraction and sequencing

Genomic DNA was extracted from four parents and 110 phenotyped individuals from the BC1 and 110 from the field populations using the modified CTAB protocol [65]. DNA concentration and quality were assessed using spectrometry (NanoDrop^®^) and 1% agarose gel electrophoresis in 1x TAE buffer. Following Elshire et al. (2011) [66], Genotyping by Sequencing (GBS) libraries were prepared using *PstI* restriction enzyme, individual tags, a common adapter, and barcodes for sample identification. Sequencing was conducted on the Illumina HiSeq 2500^®^ platform.

### Data pre-processing

Reads were demultiplexed and trimmed using Stacks (version 2.62) [67] and aligned to the *H. armigera* genome assembly GCF_030705265.1 (BioProject: PRJNA713413) [17] using BWA (version 2.0) [68]. An average alignment rate of ∼95% was achieved. SAM files were converted to BAM format using Samtools (version 1.4) [69].

SNP calling was performed using the Stacks pipeline. First, a map of each population was generated in TXT format, which was then used by subsequent commands in the Stacks program. The first column of this map contains the sample names, and the second column indicates the population to which each sample belongs. Subsequently, the gstacks command was used to assemble loci, perform variant calling, and generate libraries of loci and variants. Finally, the populations command was employed to export the data in VCF format. Data filtering was conducted using VCFtools, retaining SNPs with Genotype Quality > 40, Haplotype Quality > 20, Read Depth > 5, and a maximum of 25% missing data.

### GWAS analysis

Population structure analysis was executed using the Discriminant Analysis of Principal Components (DAPC) model from the DAPC package [70] in R software [71]. The Genome Association and Prediction Integrated Tool (GAPIT) package [72] was employed for the association study. The GAPIT function uses the HAPMAP format, so the VCF file was converted by the vcfR package [73] in R.

The GWAS utilized the Blink model within the GAPIT function, integrating parameters like the number of principal components (PCA) estimated (K) and minor allele frequency (MAF) set at 0.01. The Blink model, designed to mitigate the assumption of evenly distributed causal genes across the genome, selectively includes or excludes genes based on linkage disequilibrium (LD) signals.

SNP significance was established using the Bonferroni multiple test correction, supplemented by a permutation test with 10,000 iterations, setting a global 5% significance level for type I error. Pairwise LD around significant SNPs was calculated using the GAPIT package [72], adopting the R^2^ method with a 10-SNP moving window. The high LD region surrounding significant SNPs was scrutinized for potential high-field survival genes in the *H. armigera* genome.

### QTL mapping

The linkage map was constructed by the LepMap3 program [74]. The pedigree design used to construct the linkage maps consists of two grandparents (♂ Flub-R and ♀ TWBS) who are the parents of the heterozygous individual (♂ HET) and two dummy grandparents (♂ GP1 and ♀ GP2) who are the parents of the susceptible female (♀ TWBS2), used for backcrossing with the heterozygous (♂ HET). Additionally, it includes 109 BC1 individuals (S5 Fig). The ParentCall function was executed with the parameters removeNonInformative=1. Due to the previous filter, we did not use the Filtering2 function. The ordered arrangement of markers within each chromosome was achieved using the OrderMarkers2 function, employing the outputPhasedData=1, recombination2=0, useKosambi=1, proximityScale=100 and usePhysical=1 0.1 parameters. The OrderMarkers2 function was run individually for each chromosome, and after this, the markers ordered were converted to genotypes by the map2genotypes.awk script from LepMap3. Ultimately, the markers’ names were recovered by the script ChangeMarkerNames.awk, and the chromosomes were merged into one file.

QTL mapping for the resistance trait employed the r/QTL program [75]. Chromosomal TXT files were manually converted to CSV format, with genotypes 11, 12, 21, and 22 transcribed as AB, AB, AA, and AA, respectively (S5 Fig). The CSV file, incorporating the linkage group map, genotypes, and phenotypes, was imported into r/QTL. Data quality was ensured by removing markers with low genotypic information and merging markers at the same position. Genotype imputation was executed using the fill.geno function and genotypic probability were calculated using the calc.genoprob function.

Interval mapping (IM) analysis was performed using the scanone function, considering binomial regression for survival phenotype (binary variable). The LOD threshold for significance was determined through a permutation test with 10,000 iterations. The makeqtl and fitqtl functions analyzed LOD scores and effects for each QTL. The bayesint function estimated QTL confidence intervals while flanking markers determined the locus size in centiMorgans. An in-depth investigation identified candidate genes associated with flubendiamide resistance within the QTL interval.

### Candidate genes

To investigate potential genes associated with *H. armigera* resistance to flubendiamide, we used a 300 kb upstream and downstream of regions near each significant SNP identified. This process utilized the genome annotation file (GCF_030705265.1-RS_2024_03) from *H. armigera* genome assembly GCF_030705265.1 published in the NCBI.

## Acknowledgments

We thank the Brazilian Insecticide Resistance Action Committee (IRAC-BR) for providing *Helicoverpa armigera* populations for this research. We also thank the Statistical Genetics Laboratory - ESALQ/USP and the Insect Evolution and Genomics Group – University of Cambridge team for all support in this work.

## Data availability statement

The raw genomic sequencing data of the *Helicoverpa armigera* field populations and laboratory strains are available in the Sequence Read Archive (SRA) under the BioProject accession number PRJNA1139396.

