## Supplemental Figure 1 for "The genetic architecture of resistance to flubendiamide insecticides in *Helicoverpa armigera* (Hübner) (Lepidoptera: Noctuidae)"

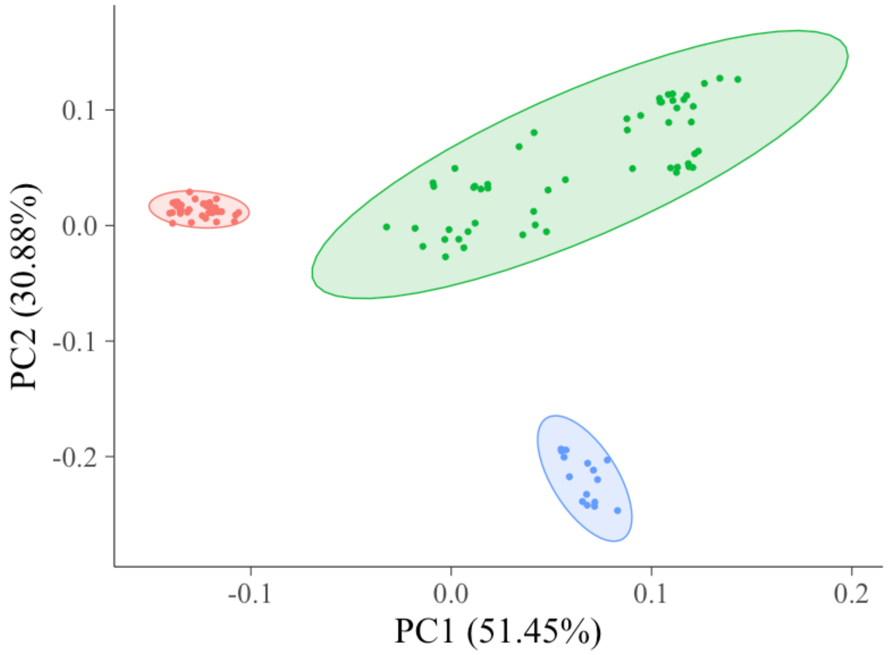


**S1 Fig. Two-dimensional discriminant principal component analysis of *Helicoverpa armigera* field population**. The colours represent the sample clusters identified by PCA analysis, indicating K = 3. The circles denote the Euclidean distance from the centre of each cluster, corresponding to the 95% confidence ellipse.
