## Supplemental Figure 5 for "The genetic architecture of resistance to flubendiamide insecticides in *Helicoverpa armigera* (Hübner) (Lepidoptera: Noctuidae)"

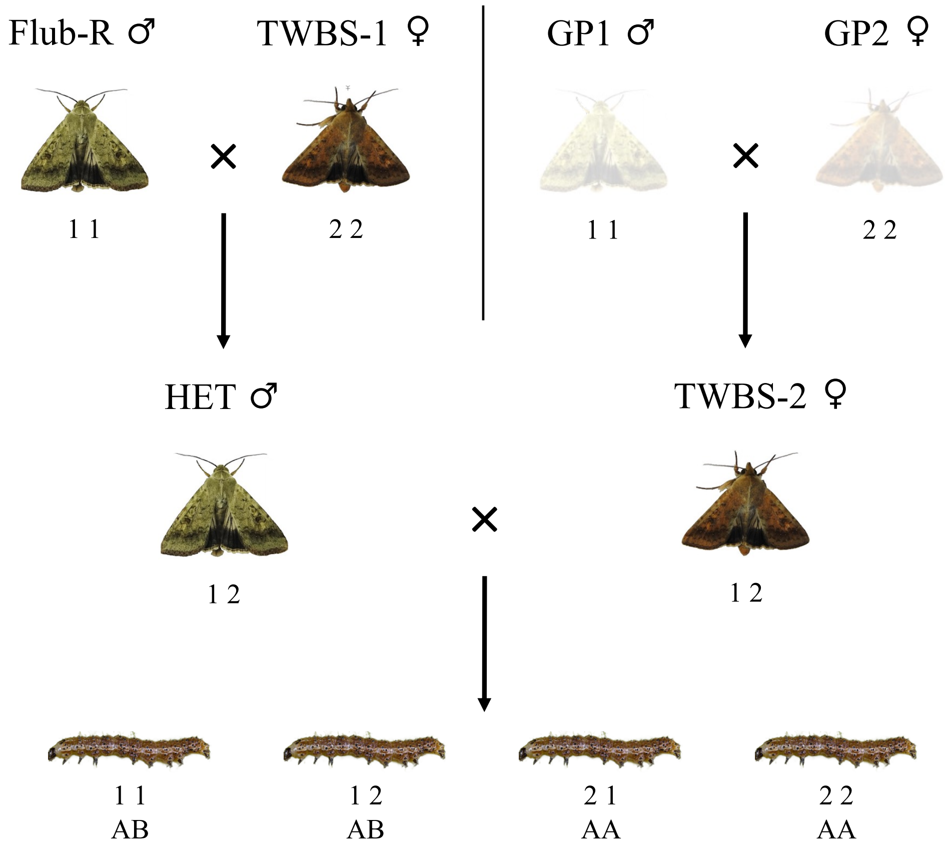


**S5 Fig. The pedigree design used in the linkage map construction by LepMap3.** Grandparents 1 and 2, shown as semi-transparent, represent dummy grandparents that were added to the pedigree file of the BC1 population, as indicated by the LepMap3 manual. The values represent the genotype codes used by the programme, where 1 1 and 2 2 denote male homozygotes and female homozygotes, respectively. The codes 1 2 and 2 1 represent heterozygotes. The first number originates from the paternal side, and the second is from the maternal side. These codes were converted to AA and AB codes used by the rQTL programme, representing susceptible homozygotes and heterozygotes, respectively.
