## Supplemental Table 1 for "The genetic architecture of resistance to flubendiamide insecticides in *Helicoverpa armigera* (Hübner) (Lepidoptera: Noctuidae)"

**S1 Table.** Linkage map summary.

| LG | No. Markers | Length (cM) | Mean Spacing (cM) | Max Spacing (cM) |
|---|---|---|---|---|
| 1 | 46 | 118.94 | 2.64 | 11.19 |
| 2 | 37 | 110.76 | 3.08 | 14.13 |
| 3 | 47 | 118.63 | 2.58 | 24.78 |
| 4 | 31 | 121.68 | 4.06 | 19.25 |
| 5 | 52 | 127.96 | 2.51 | 19.25 |
| 6 | 41 | 106.68 | 2.67 | 19.25 |
| 7 | 39 | 116.21 | 3.06 | 10.23 |
| 8 | 49 | 133.70 | 2.79 | 10.23 |
| 9 | 50 | 107.31 | 2.19 | 30.95 |
| 10 | 45 | 124.58 | 2.83 | 12.16 |
| 11 | 45 | 145.38 | 3.30 | 16.13 |
| 12 | 45 | 117.88 | 2.68 | 11.19 |
| 13 | 46 | 135.06 | 3.00 | 15.12 |
| 14 | 36 | 104.26 | 2.98 | 21.40 |
| 15 | 40 | 123.02 | 3.15 | 14.13 |
| 16 | 38 | 119.74 | 3.24 | 18.19 |
| 17 | 25 | 103.66 | 4.32 | 32.29 |
| 18 | 32 | 91.56 | 2.95 | 15.12 |
| 19 | 27 | 120.22 | 4.62 | 24.79 |
| 20 | 47 | 108.65 | 2.36 | 12.16 |
| 21 | 33 | 117.73 | 3.68 | 15.12 |
| 22 | 44 | 118.71 | 2.76 | 8.33 |
| 23 | 43 | 137.88 | 3.28 | 22.51 |
| 24 | 18 | 115.27 | 6.78 | 23.64 |
| 25 | 26 | 140.68 | 5.63 | 23.63 |
| 26 | 27 | 103.16 | 3.97 | 19.25 |
| 27 | 14 | 113.88 | 8.76 | 24.78 |
| 28 | 32 | 153.26 | 4.94 | 18.19 |
| 29 | 18 | 123.98 | 7.29 | 20.32 |
| 30 | 22 | 127.62 | 6.08 | 27.16 |
| 31 | 23 | 114.71 | 5.21 | 19.25 |
