## Supplemental Figure 2 for "The genetic architecture of resistance to flubendiamide insecticides in *Helicoverpa armigera* (Hübner) (Lepidoptera: Noctuidae)"

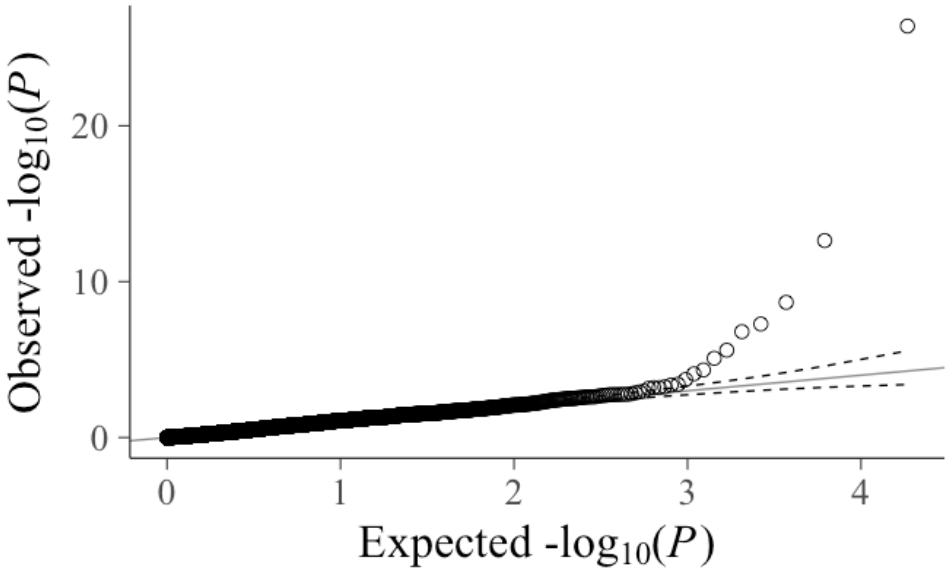


**S2 Fig. The Quantile-Quantile plot indicates the fitness of the Blink model for survival association analysis.** The light grey line shows the $-\log_{10}$(*P-values*) expected. The dashed lines represent the upper and lower limits of the 95% confidence interval. The black unfilled circles show the $-\log_{10}$(*P-values*) observed.
