## Supplemental Table 2 for "The genetic architecture of resistance to flubendiamide insecticides in *Helicoverpa armigera* (Hübner) (Lepidoptera: Noctuidae)"

**S2 Table.** List of genes within a 300 Kb downstream and upstream from the markers identified by GWAS and QTL mapping, linked to *H. armigera* survival to flubendiamide.

| Chromosome | RefSeq | Start | End | Strand | Gene ID | Description | Associated SNPs (Statistical Analysis) |
|---|---|---|---|---|---|---|---|
| Z | NC_087120.1 | 3156825 | 3168521 | + | LOC110379460 | uncharacterized protein CG1161 | |
| Z | NC_087120.1 | 3169578 | 3182738 | - | LOC110379465 | uncharacterized LOC110379465 | |
| Z | NC_087120.1 | 3183368 | 3217595 | + | LOC110379468 | roquin-1 | |
| Z | NC_087120.1 | 3224958 | 3247304 | + | LOC110379469 | uncharacterized LOC110379469 | |
| Z | NC_087120.1 | 3250802 | 3259898 | + | LOC110373007 | atypical kinase COQ8B-2C mitochondrial | |
| Z | NC_087120.1 | 3268500 | 3278214 | + | Wrm1 | wurmchen 1 | |
| Z | NC_087120.1 | 3271193 | 3277075 | - | LOC135118117 | uncharacterized LOC135118117 | |
| Z | NC_087120.1 | 3280642 | 3281665 | - | LOC110373795 | peptidyl-prolyl cis-trans isomerase-like 1 | |
| Z | NC_087120.1 | 3282086 | 3303330 | + | LOC110373753 | rho GTPase-activating protein 1 | |
| Z | NC_087120.1 | 3305424 | 3349850 | + | LOC110373799 | PHD finger protein rhinoceros | |
| Z | NC_087120.1 | 3360832 | 3475315 | + | Nwk | nervou swreck | rs1P3460302 (GWAS) |
| Z | NC_087120.1 | 3481796 | 3492756 | - | LOC110379461 | atypical kinase COQ8B-2C mitochondrial | |
| Z | NC_087120.1 | 3501037 | 3517277 | - | LOC110373023 | Phosphatidylserine lipase ABHD16A | |
| Z | NC_087120.1 | 3518186 | 3536343 | + | LOC110373032 | protein GPR107 | |
| Z | NC_087120.1 | 3552592 | 3579679 | + | LOC110373030 | ras-relatedandestrogen-regulated growth inhibitor | |
| Z | NC_087120.1 | 3583124 | 3593025 | + | LOC110373024 | nuclear receptor subfamily 2 group E member 1 | |
| Z | NC_087120.1 | 3597157 | 3649026 | + | LOC110373000 | calcium-binding mitochondrial carrier protein SCaMC-2 | |
| Z | NC_087120.1 | 3653786 | 3665298 | - | LOC110373012 | lysine-specific demethylase 3A-A | |
| Z | NC_087120.1 | 3667365 | 3683354 | + | LOC110372998 | ATP-binding cassette sub-family F member 2 | |
| Z | NC_087120.1 | 3683355 | 3902488 | - | LOC126054100 | uncharacterized LOC126054100 | |
| Z | NC_087120.1 | 3750422 | 3750495 | + | Trnai-aau | transfer RNA isoleucine (anticodon AAU) | |
| 2 | NC_087121.1 | 1634694 | 1637974 | + | LOC110376521 | lipase member H | |
| 2 | NC_087121.1 | 1660167 | 1661511 | - | LOC110376516 | lipase member H | |
| 2 | NC_087121.1 | 1663158 | 1671650 | - | LOC110376523 | lipase member I | |
| 2 | NC_087121.1 | 1666802 | 1670124 | - | LOC110376524 | lipase member H | |
| 2 | NC_087121.1 | 1690778 | 1694332 | - | LOC135118694 | uncharacterized LOC135118694 | |
| 2 | NC_087121.1 | 1701866 | 1703839 | - | LOC126055277 | uncharacterized LOC126055277 | |
| 2 | NC_087121.1 | 1713506 | 1713577 | - | Trnaa-cgc-3 | transfer RNA alanine (anticodon CGC) | |
| 2 | NC_087121.1 | 1786763 | 1788331 | + | LOC110376529 | uncharacterized LOC110376529 | |
| 2 | NC_087121.1 | 1806428 | 1820115 | + | LOC110376514 | uncharacterized LOC110376514 | |
| 2 | NC_087121.1 | 1840635 | 1941818 | + | LOC110372095 | uncharacterized LOC110372095 | rs2P1937820 (Lower boundary of the QTL) |
| 2 | NC_087121.1 | 1943506 | 1945479 | + | LOC126056517 | uncharacterized LOC126056517 | |
| 2 | NC_087121.1 | 2023799 | 2023870 | - | Trnad-guc-3 | transfer RNA aspartic acid (anticodon GUC) | |
| 2 | NC_087121.1 | 2026439 | 2027764 | + | LOC126055600 | uncharacterized LOC126055600 | |
| 2 | NC_087121.1 | 2027222 | 2028898 | - | LOC126055594 | uncharacterized LOC126055594 | |
| 2 | NC_087121.1 | 2048721 | 2059694 | - | LOC110372073 | probable 3-hydroxyisobutyrate dehydrogenase%2C mitochondrial | |
| 2 | NC_087121.1 | 2065610 | 2074390 | - | LOC110372049 | sulfotransferase 1E1 | |
| 2 | NC_087121.1 | 2092901 | 2106319 | + | LOC110372097 | sulfotransferase 1C3 | |
| 2 | NC_087121.1 | 2114207 | 2193215 | - | LOC110372027 | protein tweety | |
| 2 | NC_087121.1 | 2140556 | 2147706 | + | LOC126054659 | uncharacterized LOC126054659 | |
| 2 | NC_087121.1 | 2178498 | 2180448 | + | LOC126054004 | uncharacterized LOC126054004 | |
| 2 | NC_087121.1 | 2180586 | 2182469 | - | LOC135117689 | putative nuclease HARBI1 | |
| 2 | NC_087121.1 | 2193487 | 2195705 | + | LOC110372086 | cytosolic Fe-S cluster assembly factor Nubp2 homolog | |
| 2 | NC_087121.1 | 2196121 | 2215720 | - | LOC110372048 | cryptochrome-1 | |
| 2 | NC_087121.1 | 2215819 | 2216848 | + | LOC110372042 | uncharacterized LOC110372042 | |
| 2 | NC_087121.1 | 2218022 | 2227209 | + | LOC110372089 | uncharacterized LOC110372089 | |
| 2 | NC_087121.1 | 2227169 | 2236910 | - | LOC110372088 | uncharacterized LOC110372088 | |

| | | | | | | | |
|---|---|---|---|---|---|---|---|
| 2 | NC_087121.1 | 2458435 | 2463133 | - | LOC110372053 | developmentally-regulated GTP-binding protein 2 | |
| 2 | NC_087121.1 | 2464426 | 2465938 | + | LOC110372072 | uncharacterized LOC110372072 | |
| 2 | NC_087121.1 | 2466302 | 2471710 | + | LOC110372077 | probable RNA-binding protein 46 | |
| 2 | NC_087121.1 | 2471880 | 2480711 | + | LOC110372076 | uncharacterized LOC110372076 | |
| 2 | NC_087121.1 | 2489368 | 2565140 | - | Myo10a | Myosin 10A | |
| 2 | NC_087121.1 | 2566739 | 2567563 | + | LOC110372060 | protein BUD31 homolog | |
| 2 | NC_087121.1 | 2568025 | 2571380 | - | LOC110372059 | innexin inx2 | |
| 2 | NC_087121.1 | 2572045 | 2596108 | - | LOC110372057 | alkyldihydroxyacetonephosphate synthase | |
| 2 | NC_087121.1 | 2596839 | 2647954 | - | LOC110372051 | uncharacterized LOC110372051 | |
| 2 | NC_087121.1 | 2664567 | 2665685 | + | LOC110372085 | protein transport protein SEC31 | |
| 2 | NC_087121.1 | 2667212 | 2671132 | + | LOC110372078 | E3 ubiquitin-protein ligase RING1 | |
| 2 | NC_087121.1 | 2675090 | 2676281 | + | LOC110372079 | large ribosomal subunit protein eL22 | |
| 2 | NC_087121.1 | 2677399 | 2697202 | - | LOC110372026 | probable aconitate hydratase, mitochondrial | |
| 2 | NC_087121.1 | 2697648 | 2705986 | + | Sec15 | Secretory 15 | |
| 2 | NC_087121.1 | 2723805 | 2727015 | - | LOC110372061 | fatty acid-binding protein, liver | |
| 2 | NC_087121.1 | 2732066 | 2761175 | + | LOC110372044 | uncharacterized LOC110372044 | rs2P2759433 (GWAS), rs2P2760409 (QTL Peak) |
| 2 | NC_087121.1 | 2761147 | 2804657 | - | LOC110372045 | adenylosuccinate lyase | |
| 2 | NC_087121.1 | 2765810 | 2801220 | + | LOC110372047 | uncharacterized LOC110372047 | |
| 2 | NC_087121.1 | 2805181 | 2814541 | - | LOC110372043 | hydroxysteroid dehydrogenase-like protein 2 | |
| 2 | NC_087121.1 | 2818825 | 2884188 | - | Wake | wide awake | |
| 2 | NC_087121.1 | 2890476 | 2891159 | + | LOC135117725 | uncharacterized LOC135117725 | |
| 2 | NC_087121.1 | 2923606 | 3082866 | - | LOC135118708 | uncharacterized LOC135118708 | |
| 2 | NC_087121.1 | 2998356 | 3003561 | - | LOC135117926 | uncharacterized LOC135117926 | |
| 2 | NC_087121.1 | 3005268 | 3006966 | + | LOC110372036 | uncharacterized LOC110372036 | |
| 2 | NC_087121.1 | 3473236 | 3479522 | + | LOC110372068 | uncharacterized LOC110372068 | |
| 2 | NC_087121.1 | 3481409 | 3484180 | - | LOC110372069 | heparan sulfate glucosamine 3-O-sulfotransferase 3A1 | |
| 2 | NC_087121.1 | 3484581 | 3485938 | + | LOC110372071 | uncharacterized LOC110372071 | |
| 2 | NC_087121.1 | 3486097 | 3488533 | - | LOC110372070 | hydroxyacylglutathione hydrolase, mitochondrial | |
| 2 | NC_087121.1 | 3488650 | 3500367 | + | LOC110372067 | small subunit processome component 20 homolog | |
| 2 | NC_087121.1 | 3502410 | 3548390 | - | LOC110372075 | uncharacterized LOC110372075 | |
| 2 | NC_087121.1 | 3549545 | 3558695 | + | LOC110372052 | histone-binding protein N1/N2 | |
| 2 | NC_087121.1 | 3558978 | 3560164 | - | LOC110372083 | prostaglandin reductase 1 | |
| 2 | NC_087121.1 | 3564669 | 3565806 | - | LOC110372056 | prostaglandin reductase 1 | |
| 2 | NC_087121.1 | 3567498 | 3568599 | - | LOC110372096 | prostaglandin reductase 1 | |
| 2 | NC_087121.1 | 3571882 | 3572989 | - | LOC110375835 | prostaglandin reductase 1 | |
| 2 | NC_087121.1 | 3576958 | 3578062 | - | LOC126053453 | prostaglandin reductase 1 | |
| 2 | NC_087121.1 | 3585598 | 3586701 | - | LOC126055293 | prostaglandin reductase 1 | |
| 2 | NC_087121.1 | 3589539 | 3610946 | + | LOC110375870 | uncharacterized LOC110375870 | |
| 2 | NC_087121.1 | 3590632 | 3591773 | - | LOC135116681 | prostaglandin reductase 1-like | |
| 2 | NC_087121.1 | 3592546 | 3594011 | - | LOC110375848 | prostaglandin reductase 1 | |
| 2 | NC_087121.1 | 3594503 | 3595653 | - | LOC110375828 | prostaglandin reductase 1 | |
| 2 | NC_087121.1 | 3598844 | 3599990 | - | LOC135118795 | prostaglandin reductase 1-like | |
| 2 | NC_087121.1 | 3603509 | 3604670 | - | LOC135117746 | prostaglandin reductase 1-like | |
| 2 | NC_087121.1 | 3606513 | 3607630 | - | LOC110375859 | prostaglandin reductase 1 | |
| 2 | NC_087121.1 | 3611602 | 3619942 | - | LOC110377250 | neprilysin-4 | |
| 2 | NC_087121.1 | 3620027 | 3624803 | + | LOC110377256 | uncharacterized LOC110377256 | |
| 2 | NC_087121.1 | 3625644 | 3633453 | + | LOC110376600 | E3 ubiquitin-protein ligase CHIP | |
| 2 | NC_087121.1 | 3634549 | 3635392 | - | LOC135117752 | uncharacterized LOC135117752 | |

| | | | | | | | |
|---|---|---|---|---|---|---|---|
| 2 | NC_087121.1 | 3642556 | 3755153 | + | LOC110375503 | uncharacterized protein | |
| 2 | NC_087121.1 | 3726561 | 3728044 | + | LOC135118859 | uncharacterized LOC135118859 | |
| 2 | NC_087121.1 | 3728111 | 3731254 | + | LOC126054309 | uncharacterized LOC126054309 | |
| 2 | NC_087121.1 | 3756360 | 3775657 | - | LOC110376418 | protein CLEC16A homolog | |
| 2 | NC_087121.1 | 3777311 | 3864707 | - | LOC110377735 | uncharacterized LOC110377735 | rs2P3779183 (GWAS) |
| 2 | NC_087121.1 | 3877409 | 3879356 | - | LOC110377162 | uncharacterized LOC110377162 | |
| 2 | NC_087121.1 | 3880914 | 3884237 | + | LOC110376954 | apyrase | |
| 2 | NC_087121.1 | 3884593 | 3896675 | - | LOC110376365 | mitochondrial glycine transporter A | |
| 2 | NC_087121.1 | 3900453 | 3911255 | - | LOC110375876 | myophilin | |
| 2 | NC_087121.1 | 3914760 | 3915758 | - | LOC110376754 | uncharacterized LOC110376754 | |
| 2 | NC_087121.1 | 3916487 | 3918276 | - | LOC110377188 | uncharacterized LOC110377188 | |
| 2 | NC_087121.1 | 3918961 | 3924593 | - | LOC110376877 | uncharacterized LOC110376877 | |
| 2 | NC_087121.1 | 3926571 | 3928731 | + | LOC110376964 | alpha-(1%2C3)-fucosyltransferase C | |
| 2 | NC_087121.1 | 3928922 | 3930605 | - | LOC110376868 | alpha-(1%2C3)-fucosyltransferase C | |
| 2 | NC_087121.1 | 3932414 | 3933628 | - | LOC126056580 | uncharacterized LOC126056580 | |
| 2 | NC_087121.1 | 3936031 | 3938034 | + | LOC126055286 | alpha-(1%2C3)-fucosyltransferase C | |
| 2 | NC_087121.1 | 3940695 | 3942903 | + | LOC135116619 | alpha-(1%2C3)-fucosyltransferase C-like | |
| 2 | NC_087121.1 | 3944195 | 3946368 | + | LOC110377087 | alpha-(1%2C3)-fucosyltransferase C | |
| 2 | NC_087121.1 | 3946574 | 3949869 | + | LOC110377096 | alpha-(1%2C3)-fucosyltransferase C | |
| 2 | NC_087121.1 | 3963051 | 3965022 | + | LOC110377152 | uncharacterized LOC110377152 | |
| 2 | NC_087121.1 | 3965236 | 3966796 | + | LOC110375483 | uncharacterized LOC110375483 | |
| 2 | NC_087121.1 | 3967009 | 3968680 | - | LOC110375478 | uncharacterized LOC110375478 | |
| 2 | NC_087121.1 | 3970104 | 3973435 | - | LOC110375471 | uncharacterized LOC110375471 | |
| 2 | NC_087121.1 | 3992033 | 3994408 | + | LOC126055971 | uncharacterized LOC126055971 | |
| 2 | NC_087121.1 | 4002116 | 4149758 | + | LOC110375954 | voltage-dependent T-type calcium channel subunit alpha-1G | |
| 2 | NC_087121.1 | 6627440 | 6632121 | + | LOC135117972 | uncharacterized LOC135117972 | |
| 2 | NC_087121.1 | 6633716 | 6642064 | + | LOC110376138 | diacylglycerol kinase epsilon | |
| 2 | NC_087121.1 | 6642622 | 6651106 | - | LOC110376276 | mitochondrial dicarboxylate carrier | |
| 2 | NC_087121.1 | 6649684 | 6651970 | + | Elp6 | Elongator complex protein 6 | |
| 2 | NC_087121.1 | 6653206 | 6671915 | + | LOC110376105 | kinesin-like protein KIF19 | |
| 2 | NC_087121.1 | 6672231 | 6688522 | - | LOC110376163 | pH-sensitive chloride channel 2 | |
| 2 | NC_087121.1 | 6688382 | 6691678 | - | LOC110376095 | transducin beta-like protein 3 | |
| 2 | NC_087121.1 | 6691795 | 6692946 | + | LOC110376340 | transmembrane protein 186 | |
| 2 | NC_087121.1 | 6693044 | 6696010 | + | LOC135117988 | uncharacterized LOC135117988 | |
| 2 | NC_087121.1 | 6696024 | 6697336 | - | LOC135117984 | putative nuclease HARBI1 | |
| 2 | NC_087121.1 | 6697621 | 6701597 | - | LOC110376193 | dual specificity mitogen-activated protein kinase kinase 4 | |
| 2 | NC_087121.1 | 6702136 | 6704686 | + | LOC110376267 | DNA-directed RNA polymerase I subunit RPA43 | |
| 2 | NC_087121.1 | 6706824 | 6708327 | + | LOC110376302 | small ribosomal subunit protein uS7m | |
| 2 | NC_087121.1 | 6715967 | 6741543 | + | LOC110375993 | phospholipid-transporting ATPase ABCA3 | |
| 2 | NC_087121.1 | 6741170 | 6764709 | - | LOC110376003 | phospholipid-transporting ATPase ABCA1 | |
| 2 | NC_087121.1 | 6766869 | 6776563 | + | LOC110375647 | ubiquitin carboxyl-terminal hydrolase 31 | |
| 2 | NC_087121.1 | 6778146 | 6783101 | - | LOC110377503 | uncharacterized LOC110377503 | |
| 2 | NC_087121.1 | 6783647 | 6819922 | + | LOC110375548 | protein yellow | |
| 2 | NC_087121.1 | 6823287 | 6841149 | + | LOC110375518 | protein yellow | |
| 2 | NC_087121.1 | 6841269 | 6843221 | - | LOC110375358 | ATP synthase subunit O, mitochondrial | |
| 2 | NC_087121.1 | 6843847 | 6849475 | - | LOC110375580 | UPF0764 protein C16orf89 homolog | |
| 2 | NC_087121.1 | 6850173 | 6858040 | + | LOC110377298 | protein yellow | |
| 2 | NC_087121.1 | 6858580 | 6861839 | + | LOC110377309 | protein yellow | |
| 2 | NC_087121.1 | 6862124 | 6865299 | + | LOC110375349 | protein yellow | |

| | | | | | | | |
|---|---|---|---|---|---|---|---|
| 2 | NC_087121.1 | 6865424 | 6873166 | - | LOC110375339 | uncharacterized LOC110375339 | |
| 2 | NC_087121.1 | 6873554 | 6906013 | + | LOC110376662 | uncharacterized LOC110376662 | |
| 2 | NC_087121.1 | 6894050 | 6895465 | - | LOC135118013 | uncharacterized LOC135118013 | |
| 2 | NC_087121.1 | 6906619 | 6908906 | + | LOC110377132 | L-xylulose reductase | |
| 2 | NC_087121.1 | 6909347 | 6910817 | + | LOC110377141 | D-erythrulose reductase | |
| 2 | NC_087121.1 | 6910821 | 6919174 | - | LOC110377122 | casein kinase I | |
| 2 | NC_087121.1 | 6920527 | 6936164 | - | LOC110376783 | class A basic helix-loop-helix protein 15 | rs2P6931787 (GWAS) |
| 2 | NC_087121.1 | 6939491 | 6950267 | + | LOC135118004 | uncharacterized LOC135118004 | |
| 2 | NC_087121.1 | 6950268 | 6952564 | - | LOC110377015 | uncharacterized LOC110377015 | |
| 2 | NC_087121.1 | 6953213 | 6955408 | + | LOC110375774 | uncharacterized LOC110375774 | |
| 2 | NC_087121.1 | 6955182 | 6957571 | - | LOC110375765 | uncharacterized protein | |
| 2 | NC_087121.1 | 6982067 | 6996873 | + | LOC110377192 | uncharacterized LOC110377192 | |
| 2 | NC_087121.1 | 7031730 | 7033701 | + | LOC110375944 | ribosomal L1 domain-containing protein 1 | |
| 2 | NC_087121.1 | 7033612 | 7074133 | - | Nbs | nbs | |
| 2 | NC_087121.1 | 7036413 | 7048435 | + | Rempa | reduced mechanoreceptor potential A | |
| 2 | NC_087121.1 | 7044142 | 7050355 | - | LOC110377230 | uncharacterized LOC110377230 | |
| 2 | NC_087121.1 | 7046617 | 7050185 | - | LOC126056560 | uncharacterized LOC126056560 | |
| 2 | NC_087121.1 | 7050727 | 7055693 | + | LOC110375709 | translation factor GUF1 homolog, mitochondrial | |
| 2 | NC_087121.1 | 7055914 | 7060358 | - | LOC110375720 | WD repeat domain phosphoinositide-interacting protein 3 | |
| 2 | NC_087121.1 | 7061101 | 7061853 | - | LOC110376651 | ragulator complex protein LAMTOR2 homolog | |
| 2 | NC_087121.1 | 7063283 | 7065678 | - | LOC110376535 | protein SDA1 homolog | |
| 2 | NC_087121.1 | 7066134 | 7069778 | + | Kdelr | KDEL receptor | |
| 2 | NC_087121.1 | 7070997 | 7073468 | - | LOC110377236 | DNA polymerase subunit gamma-2, mitochondrial | |
| 2 | NC_087121.1 | 7079923 | 7086476 | + | LOC110377421 | aldo-keto reductase family 1 member B1 | |
| 2 | NC_087121.1 | 7087937 | 7092530 | + | LOC135118725 | uncharacterized LOC135118725 | |
| 2 | NC_087121.1 | 7096825 | 7098942 | + | LOC110383041 | putative nuclease HARBI1 | |
| 2 | NC_087121.1 | 7098877 | 7100974 | - | LOC110383055 | uncharacterized LOC110383055 | |
| 2 | NC_087121.1 | 7106324 | 7112865 | + | LOC135118029 | aldo-keto reductase family 1 member B1-like | |
| 2 | NC_087121.1 | 7114449 | 7161492 | + | LOC135118726 | uncharacterized LOC135118726 | |
| 2 | NC_087121.1 | 7182091 | 7189907 | + | LOC135118033 | uncharacterized LOC135118033 | |
| 2 | NC_087121.1 | 7190014 | 7207134 | - | LOC110377596 | chaoptin | |
| 2 | NC_087121.1 | 13805097 | 13805564 | - | LOC126055221 | uncharacterized LOC126055221 | |
| 2 | NC_087121.1 | 13848977 | 13850417 | + | LOC126055232 | uncharacterized LOC126055232 | |
| 2 | NC_087121.1 | 13851523 | 13852393 | - | LOC135118818 | uncharacterized LOC135118818 | |
| 2 | NC_087121.1 | 13871344 | 13913818 | + | LOC135116632 | lethal(2) giant larvae protein homolog 1-like | |
| 2 | NC_087121.1 | 13916795 | 13922310 | + | LOC135118821 | oocyte zinc finger protein XlCOF6.1-like | |
| 2 | NC_087121.1 | 13923168 | 13930539 | - | LOC135118822 | coiled-coil domain-containing protein 40-like | |
| 2 | NC_087121.1 | 13930526 | 13935849 | + | LOC135118766 | intraflagellar transport protein 56-like | |
| 2 | NC_087121.1 | 13938182 | 13940505 | - | LOC135118884 | cuticle protein 21-like | |
| 2 | NC_087121.1 | 13943292 | 13946723 | - | LOC110379225 | sodium/potassium-transporting ATPase subunit alpha | |
| 2 | NC_087121.1 | 13949681 | 13964220 | - | LOC135118493 | WD repeat-containing protein 81-like | |
| 2 | NC_087121.1 | 13964580 | 13966789 | - | LOC135118496 | uridine 5'-monophosphate synthase-like | |
| 2 | NC_087121.1 | 13974042 | 13996862 | + | LOC135118767 | zinc finger protein 436-like | |
| 2 | NC_087121.1 | 13998256 | 14004246 | - | LOC110379415 | zinc finger protein 689 | |
| 2 | NC_087121.1 | 14009513 | 14020390 | - | LOC110378434 | zinc finger protein with KRAB and SCAN domains 7 | |
| 2 | NC_087121.1 | 14061235 | 14062661 | - | LOC135118505 | uncharacterized LOC135118505 | |
| 2 | NC_087121.1 | 14062827 | 14065091 | + | LOC110378449 | actin-5C | |
| 2 | NC_087121.1 | 14066584 | 14095088 | - | LOC110377798 | klaroid protein | |
| 2 | NC_087121.1 | 14095427 | 14116964 | + | LOC135118502 | Fanconi anemia group I protein homolog | rs2P14104337 (Upper boundary of the QTL) |

| | | | | | | | |
|---|---|---|---|---|---|---|---|
| 2 | NC_087121.1 | 14130057 | 14130874 | + | LOC135118389 | uncharacterized LOC135118389 | |
| 2 | NC_087121.1 | 14148376 | 14219924 | - | LOC135118768 | uncharacterized LOC135118768 | |
| 2 | NC_087121.1 | 14180157 | 14180786 | - | LOC135118507 | uncharacterized LOC135118507 | |
| 2 | NC_087121.1 | 14224577 | 14226272 | + | LOC135118508 | NADH dehydrogenase [ubiquinone] 1 beta subcomplex subunit 10-like | |
| 2 | NC_087121.1 | 14228103 | 14245459 | - | LOC135118510 | meiosis regulator and mRNA stability factor 1-like | |
| 2 | NC_087121.1 | 14246107 | 14256116 | + | LOC135118511 | 26S proteasome non-ATPase regulatory subunit 12-like | |
| 2 | NC_087121.1 | 14256507 | 14260129 | + | LOC110375695 | uncharacterized LOC110375695 | |
| 2 | NC_087121.1 | 14260223 | 14270956 | - | LOC135118513 | inositol polyphosphate 5-phosphatase E-like | |
| 2 | NC_087121.1 | 14273372 | 14280241 | - | LOC135118514 | uncharacterized LOC135118514 | |
| 2 | NC_087121.1 | 14283177 | 14316851 | + | LOC135116537 | putative inorganic phosphate cotransporter | |
| 2 | NC_087121.1 | 14316842 | 14327874 | - | LOC110377209 | RAD50-interacting protein 1 | |
| 2 | NC_087121.1 | 14328104 | 14332895 | + | LOC135118524 | lysophospholipid acyltransferase 7-like | |
| 2 | NC_087121.1 | 14332687 | 14348594 | - | LOC110377700 | keratin%2C type I cytoskeletal 9 | |
| 13 | NC_087132.1 | 3867903 | 3872274 | - | LOC110384348 | tyrosine-protein kinase transmembrane receptor Ror | |
| 13 | NC_087132.1 | 3873100 | 3874636 | - | LOC110384349 | thyrostimulin beta-5 subunit | |
| 13 | NC_087132.1 | 3878673 | 3880298 | + | LOC110384456 | uncharacterized LOC110384456 | |
| 13 | NC_087132.1 | 3880709 | 3892911 | + | LOC110384457 | leucine-rich repeat-containing protein 40 | |
| 13 | NC_087132.1 | 3902663 | 3920466 | + | LOC110384403 | uncharacterized LOC110384403 | |
| 13 | NC_087132.1 | 3921226 | 3923037 | - | LOC110384417 | pro-resilin | |
| 13 | NC_087132.1 | 3923709 | 3925558 | + | LOC110384401 | protein SCO1 homolog, mitochondrial | |
| 13 | NC_087132.1 | 3925686 | 3930172 | - | LOC135117704 | zinc finger protein 660-like | |
| 13 | NC_087132.1 | 3930215 | 3935858 | - | LOC110384400 | zinc finger protein 112 | |
| 13 | NC_087132.1 | 3939433 | 3940993 | + | LOC110384384 | mitochondrial basic amino acids transporter | |
| 13 | NC_087132.1 | 3941409 | 3951540 | - | LOC110384382 | kelch-like protein 10 | |
| 13 | NC_087132.1 | 3951663 | 3966400 | - | Lkrsdh | Lysine ketoglutarate reductase/saccharopine dehydrogenase | |
| 13 | NC_087132.1 | 3956460 | 3956531 | + | Trnae-uuc-5 | transfer RNA glutamic acid (anticodon UUC) | |
| 13 | NC_087132.1 | 3968177 | 3988811 | + | LOC110384363 | ethanolaminephosphotransferase 1 | |
| 13 | NC_087132.1 | 3990213 | 4122492 | - | LOC110384364 | RNA polymerase II transcriptional coactivator | |
| 13 | NC_087132.1 | 3991445 | 3993013 | + | LOC110384344 | uncharacterized LOC110384344 | |
| 13 | NC_087132.1 | 3992985 | 4003965 | - | LOC110384342 | formin-2 | |
| 13 | NC_087132.1 | 4014340 | 4038780 | + | LOC110384404 | uncharacterized LOC110384404 | |
| 13 | NC_087132.1 | 4046670 | 4101214 | - | LOC110384339 | hemicentin-2 | |
| 13 | NC_087132.1 | 4142457 | 4154672 | + | LOC110384365 | esterase FE4 | |
| 13 | NC_087132.1 | 4156145 | 4158871 | - | LOC110384366 | splicing factor U2af 38 kDa subunit | |
| 13 | NC_087132.1 | 4159329 | 4178569 | + | LOC110384360 | multidrug resistance-associated protein 1 | rs13P4171996 (GWAS) |
| 13 | NC_087132.1 | 4160534 | 4162466 | + | LOC126055707 | uncharacterized LOC126055707 | |
| 13 | NC_087132.1 | 4180513 | 4216417 | + | LOC110384372 | multidrug resistance-associated protein 1 | |
| 13 | NC_087132.1 | 4185164 | 4187608 | + | LOC126054123 | uncharacterized LOC126054123 | |
| 13 | NC_087132.1 | 4187552 | 4190243 | - | LOC126053835 | putative nuclease HARBI1 | |
| 13 | NC_087132.1 | 4217425 | 4219118 | + | LOC135117737 | uncharacterized LOC135117737 | |
| 13 | NC_087132.1 | 4222317 | 4241801 | - | LOC110384329 | A-kinase anchor protein 200 | |
| 13 | NC_087132.1 | 4242968 | 4249693 | + | LOC110384358 | transcription factor AP-4 | |
| 13 | NC_087132.1 | 4253163 | 4259663 | + | LOC110384443 | serpin B8 | |
| 13 | NC_087132.1 | 4259539 | 4260805 | - | LOC110384445 | male-enhanced antigen 1 | |
| 13 | NC_087132.1 | 4262496 | 4265223 | + | Pcmt | Protein-L-isoaspartate (D-aspartate) O-methyltransferase | |
| 13 | NC_087132.1 | 4265133 | 4269762 | - | LOC110384442 | DNA repair endonuclease XPF | |
| 13 | NC_087132.1 | 4271626 | 4289596 | + | LOC110384357 | hrp65 protein | |
| 13 | NC_087132.1 | 4290454 | 4294932 | - | LOC110371640 | lipase member H | |
| 13 | NC_087132.1 | 4296360 | 4352480 | - | LOC110371612 | protein turtle | |

| | | | | | | | |
|---|---|---|---|---|---|---|---|
| 13 | NC_087132.1 | 4355561 | 4363349 | + | LOC110371642 | polyphosphoinositide phosphatase | |
| 13 | NC_087132.1 | 4365817 | 4372347 | + | LOC110371599 | synaptojanin-1 | |
| 13 | NC_087132.1 | 4373529 | 4377186 | - | LOC110371665 | cytochrome c oxidase subunit 7A-related protein, mitochondrial | |
| 13 | NC_087132.1 | 4387966 | 4395393 | - | LOC110371623 | polyubiquitin | |
| 13 | NC_087132.1 | 4395852 | 4396899 | + | LOC110371622 | ubiquitin-ribosomal protein eL40 fusion protein | |
| 13 | NC_087132.1 | 4397837 | 4405650 | - | LOC110371621 | WD repeat-containing and planar cell polarity effector protein fritz homolog | |
| 13 | NC_087132.1 | 4424151 | 4512064 | + | LOC110371637 | pyrokinin-1 receptor | |
| 13 | NC_087132.1 | 6386520 | 6406627 | - | LOC110376158 | uncharacterized LOC110376158 | |
| 13 | NC_087132.1 | 6415823 | 6421755 | - | LOC110376146 | lipase member H | |
| 13 | NC_087132.1 | 6426243 | 6494802 | - | LOC110376155 | uncharacterized LOC110376155 | |
| 13 | NC_087132.1 | 6495673 | 6498324 | + | LOC110370743 | palmitoyltransferase ZDHHC23-A | |
| 13 | NC_087132.1 | 6498206 | 6504116 | - | LOC110370718 | girdin | |
| 13 | NC_087132.1 | 6504312 | 6505855 | + | LOC110370687 | post-GPI attachment to proteins factor 2 | |
| 13 | NC_087132.1 | 6505326 | 6509046 | - | LOC110370685 | L-2-hydroxyglutarate dehydrogenase, mitochondrial | |
| 13 | NC_087132.1 | 6509508 | 6511844 | + | LOC110370686 | modifier of mdg4 | |
| 13 | NC_087132.1 | 6512563 | 6516472 | + | LOC135117716 | transcription activator GAGA-like | |
| 13 | NC_087132.1 | 6519760 | 6527828 | + | LOC110370673 | aminoacylase-1 | |
| 13 | NC_087132.1 | 6529966 | 6542161 | + | LOC110370749 | aminoacylase-1 | |
| 13 | NC_087132.1 | 6547003 | 6569086 | + | LOC110370671 | protein dissatisfaction | |
| 13 | NC_087132.1 | 6569697 | 6571947 | - | LOC110370737 | cysteine and histidine-rich domain-containing protein morgana | |
| 13 | NC_087132.1 | 6572481 | 6601142 | + | LOC110370670 | WD repeat and FYVE domain-containing protein 3 | |
| 13 | NC_087132.1 | 6601312 | 6625598 | - | LOC110370669 | protein numb | |
| 13 | NC_087132.1 | 6625622 | 6635698 | + | LOC110370752 | xanthine dehydrogenase | |
| 13 | NC_087132.1 | 6626514 | 6628305 | + | LOC110370753 | cytochrome c oxidase assembly protein COX20, mitochondrial | |
| 13 | NC_087132.1 | 6630578 | 6630650 | + | Trnak-cuu-7 | transfer RNA lysine (anticodon CUU) | |
| 13 | NC_087132.1 | 6637884 | 6645891 | + | LOC110370678 | xanthine dehydrogenase | |
| 13 | NC_087132.1 | 6645966 | 6655706 | - | Tango1 | Transport and Golgi organization 1 | |
| 13 | NC_087132.1 | 6656475 | 6661979 | + | LOC110370751 | uncharacterized LOC110370751 | |
| 13 | NC_087132.1 | 6700310 | 6784224 | + | LOC110370738 | protein O-mannosyl-transferase TMTC2 | rs13P6678940 (GWAS) |
