## Supplemental Figure 3 for "The genetic architecture of resistance to flubendiamide insecticides in *Helicoverpa armigera* (Hübner) (Lepidoptera: Noctuidae)"

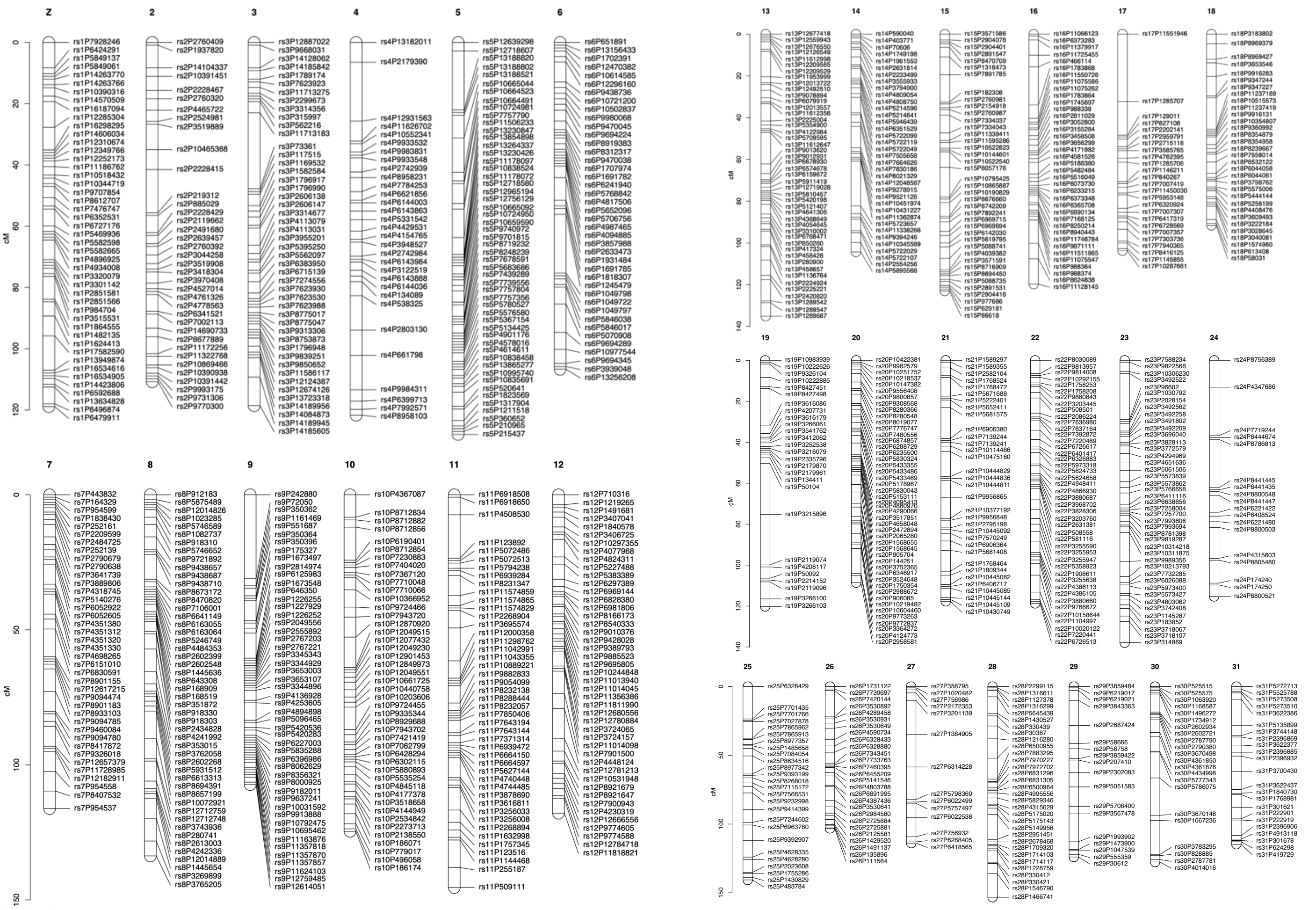

**S3 Fig. Linkage map of a *Helicoverpa armigera* backcross population, derived from the cross between the susceptible strain (TWBS) and the flubendiamide-resistant strain (Flub-R).** Each line in a linkage group denotes the position of a marker, with its respective name beside it. The y-axis shows the genetic positions of the markers and the total size of the linkage groups in centiMorgans (cM).
