## Supplemental Figure 4 for "The genetic architecture of resistance to flubendiamide insecticides in *Helicoverpa armigera* (Hübner) (Lepidoptera: Noctuidae)"

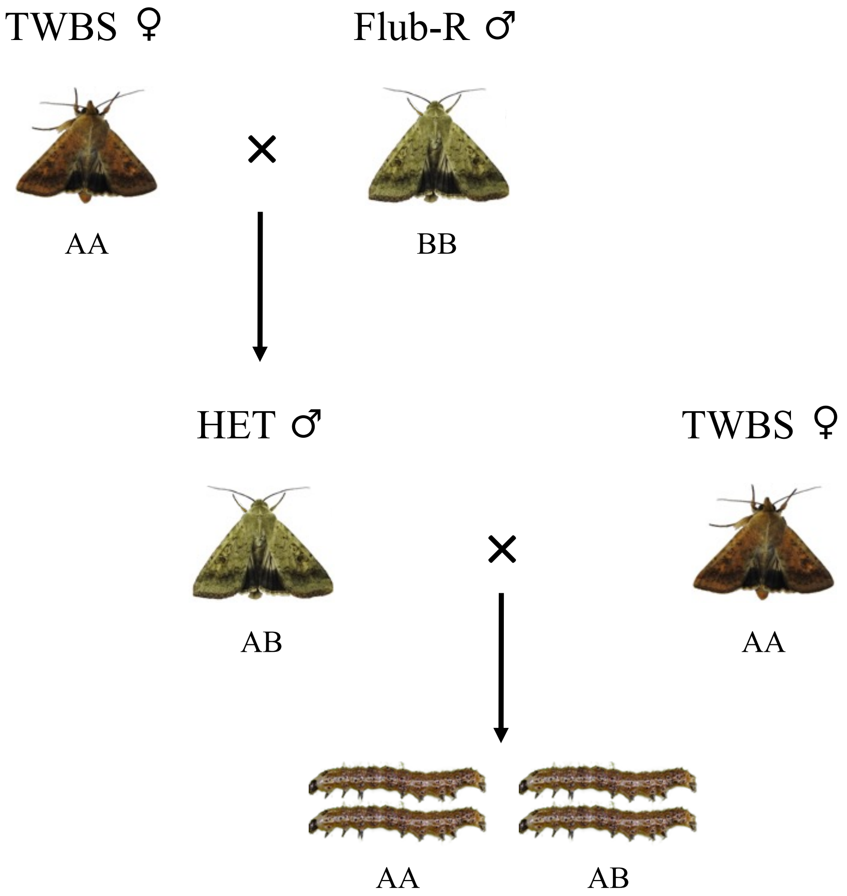


**S4 Fig. QTL mapping cross-design.** The backcross population originated from the cross between the *Helicoverpa armigera* strains Flub-R (Resistant) and TWBS (Susceptible). The larvae represent the individuals used for DNA sequencing using the GBS method. The AA code denotes the homozygous susceptible, the BB homozygous resistant and AB the heterozygous.
